# Circular-Arc Modal Migration for Ultrasound Imaging—Part II: Simulation and Experimental Evaluation in Compact Semicircular and Convex Arrays

**DOI:** 10.64898/2026.11.11.766019

**Authors:** Zihan Huang, Zhoufeng Liu, Chuan Chen, Haixia Qi, Dawei Wu

## Abstract

Circular-arc ultrasound imaging requires reconstruction that accounts for curvature and offset transmit sources across a wide field. This study evaluates how Circular-Arc Modal Migration (CAMM) changes point-response width, image-energy distribution, and low-echo target separation relative to delay-and-sum (DAS). CAMM combines angular modal analysis, local spectral remapping, and radial synthesis, with the offset phase evaluated through phase screens or approximated by scalar pullback. Guided by Part I’s error analysis and point-target comparisons, paired reconstructions use scalar pullback for compact semicircular cyst simulations and phase screens for acquisition-matched C5-2v simulations, six phantom records, and two carotid views from one participant. Each DAS/CAMM pair uses identical RF. The principal observation is that lower energy fractions outside a fixed neighborhood can coexist with broader tangential responses. In C5-2v point simulations, the full-field fraction is lower at 23 of 24 locations, whereas tangential width is greater at all 24. Semicircular cyst contrast improves in most configurations, with reversed ranking at the smallest target. All eight C5-2v cyst configurations at 10–15-mm radial depth favor CAMM in the three contrast measures; deeper comparisons depend on location and metric. Measured reflectors show the same width–energy-fraction pattern, and the evaluated phantom cyst and carotid lumen regions show improved contrast. At 512 × 513 pixels and 24 transmissions, median host latency is 29.79 ms for DAS and 51.87 ms for CAMM on an RTX 4090. The tested CAMM implementation therefore provides regional gains in low-echo target separation alongside broader tangential responses and higher computational cost.

## I. Introduction

**W** IDE-FIELD ultrasound imaging combines broad in-sonification with coherent reconstruction to recover spatial detail over an extended region. Synthetic-aperture methods achieve transmit and receive focusing through the joint processing of successive acquisitions [1]. Virtual-source formulations describe these acquisitions using a physical transmit focus as a virtual element [2], [3]. Coherent plane-wave compounding exploits angular diversity [4], while diverging transmissions provide expanding wavefronts for sector coverage [5], [6]. For circular-arc arrays, reconstruction must account for curvature and source-dependent propagation across this field.

Image quality depends on both transmit sampling and reconstruction. For convex arrays, source arrangements yielding narrow mainlobes can also produce strong off-axis responses and reduced contrast [7]. A three-dimensional diverging-wave study likewise identifies source-distribution-dependent trade-offs between resolution and secondary lobes [8]. Recent work addresses compound weights [9] and joint optimization of source positions for resolution and sidelobe performance [10]. These findings motivate evaluation across depth and angle, target size, and transmit count.

For a fixed acquisition, comparisons of virtual-source and element-based beamforming relate reconstruction to the transmitted wave and its spatial support [11]. Delay-and-sum (DAS) provides a geometrical reference whose resolution and contrast depend on receive aperture and assumed sound speed [12]. Spectral alternatives include Stolt migration for plane-wave imaging [13] and polar angular-spectrum beamforming for curvilinear arrays [14]. Circular-Arc Modal Migration (CAMM) is a direct Stolt-type construction using angular modal analysis, local radial spectral remapping, and image synthesis. Offset-source contributions are evaluated through phase-screen-weighted fields or approximated by scalar pull-back. Part I develops the common modal construction and uses theoretical error analysis and point-target comparisons to guide the offset realization [15]. Here, scalar pullback is used for the compact semicircular array and phase screens for the C5-2v convex array, linking that choice to paired simulations and recorded acquisitions. The central question is how this organization of reconstruction affects the central response, echoes outside it, and separation of low-echo targets.

A narrow point response and a clearly visible low-echo region describe different aspects of image quality. On the same RF data, a reconstruction can produce a broader central peak while showing a darker cyst interior. To characterize both aspects, we examine response width together with the fraction of image power outside a prescribed neighborhood, then assess cyst and lumen contrast using fixed paired regions. These complementary measures locate changes in response shape and target separation across depth, angle, target size, and transmit sampling.

Channel-level research systems permit different reconstructions of the same measurements [16], [17]. Shared evaluation resources such as PICMUS [18] and the UltraSound Tool- Box [19] support explicit acquisition and comparison protocols; software-defined systems such as SUPRA also assess execution cost alongside image quality [20]. Here, paired DAS/CAMM reconstructions use the same RF. Acquisition-matched simulations reproduce the convex probe geometry, firing sequence, and pulse model, connecting known-target tests to the measured-data study.

The evaluation begins with controlled cyst simulations using a compact outward-looking semicircular array with a radius of 0.75 mm and a nominal angular aperture of 180°. Cyst position, size, and transmit count are varied under this geometry. Acquisition-matched convex-array simulations then examine point and cyst responses across depth and angle using the C5-2v geometry and firing sequence of the experiments. Phantom experiments assess measured reflector responses and low-echo inclusions, followed by in vivo experiments comparing selected lumen regions in two carotid views. Finally, repeated-frame measurements quantify reconstruction latency and storage on the same workstation. Together, the simulations and experiments evaluate how the CAMM implementations change spatial response and low-echo target separation, and the computational cost of those changes. Section II presents the reconstruction workflow, and Section III specifies the acquisition, reconstruction, and evaluation protocols. Section IV reports simulation, experimental, and computational results, followed by discussion and conclusions.

## II. CAMM Reconstruction and Implementation

### A. From Channel RF to a Complex Image

CAMM reconstructs an image by reorganizing channel echoes into angular modes, remapping their spectra for radial synthesis, and evaluating the resulting fields with an offset-transmit correction. Figure 1 connects these operations to intermediate spectra and images. Its upper row contains quantities prepared from the acquisition and image grid; the lower row contains the operations repeated for each RF frame. Part I’s error analysis and point-target comparisons guide the choice of offset correction [15]. In its compact semicircular example, scalar pullback closely preserved the principal peak positions and widths obtained with phase screens. In its shallow convex example, phase screens gave lower outside-neighborhood energy fractions than scalar pullback. Guided by these results, all semicircular cyst and transmit-count studies here use scalar pullback, and all C5-2v point, cyst, phantom, and carotid reconstructions use phase screens. Part I’s Supplementary Table S1 provides starting points for other acquisition configurations.

**Fig. 1.**
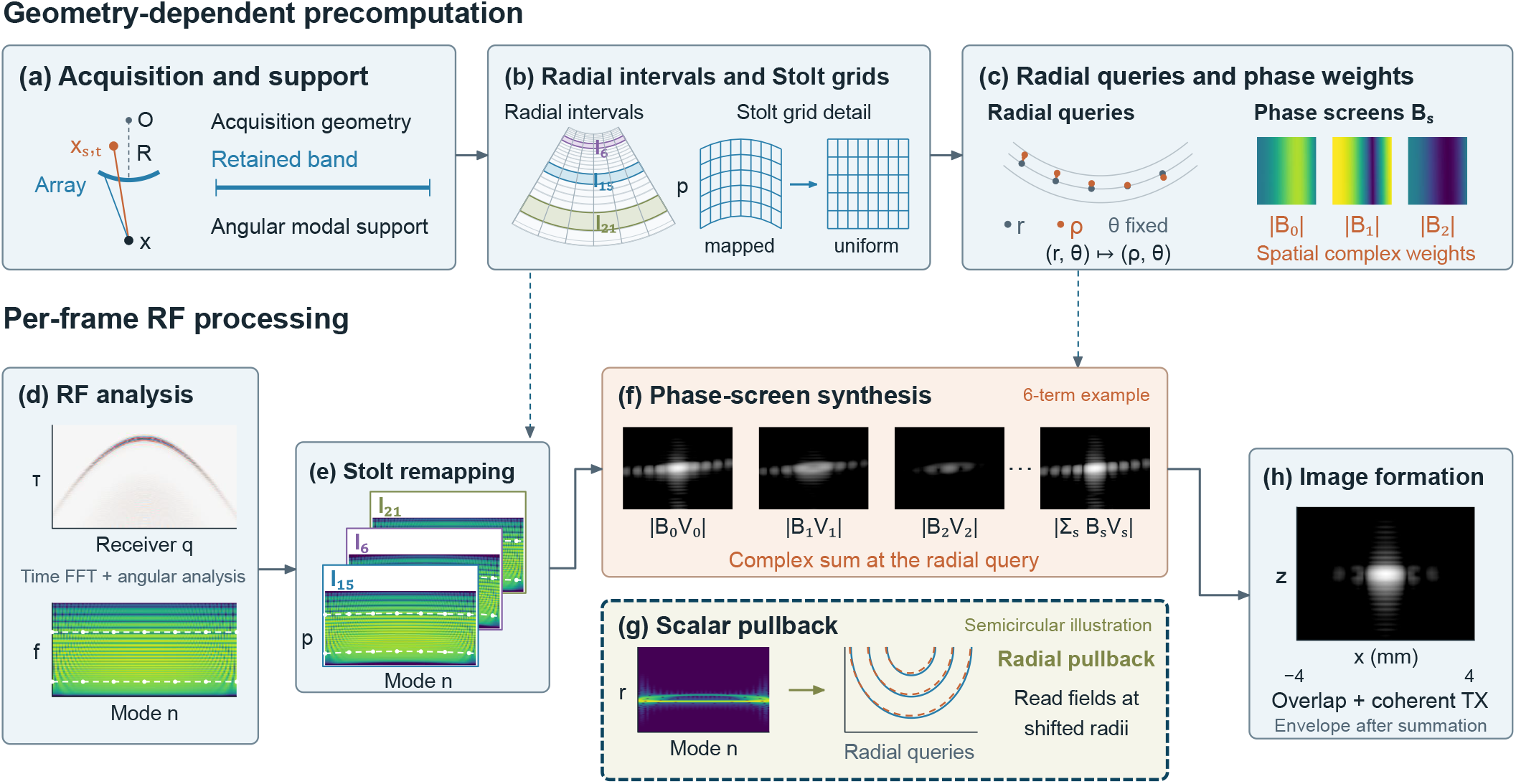
CAMM implementation. Panels (a)–(f) and (h) follow one C5-2v phase-screen example; (g) is a separate semicircular scalar-pullback illustration. Upper row: reusable geometry-dependent plans. Lower row: per-frame RF analysis, Stolt remapping, field synthesis, and coherent image formation. White traces in (d,e) track fixed input frequencies. Panel (f) displays individual contribution magnitudes alongside the magnitude of their coherent complex sum.

The input contains one time trace per receive element and transmit event. After temporal Fourier transformation, band selection, and time-origin correction, let *d*_*t*_(*k, β*_*q*_) denote the spectrum for transmission *t* and receiver angle *β*_*q*_, with acoustic wavenumber *k* = 2*πf/c*_0_ and propagation sound speed *c*_0_. Angular analysis gives

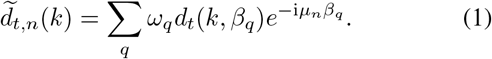

Here *ω*_*q*_ includes angular quadrature and prescribed receive weights, and *µ*_*n*_ = 2*πn/*Θ_ap_ for computational angular period Θ_ap_. Each retained *n* describes a different scale of angular variation [Fig. 1(d)].

Local Stolt remapping rearranges each modal spectrum onto a radial-wavenumber grid. For an interval centered at radius *r*_*j*_, the map is

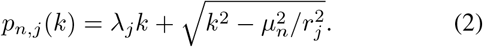

The coefficient *λ*_*j*_ is the local slope of the reference transmit path; the square-root term is the receive radial wavenumber. Only the retained propagating support is used. Interpolation onto *p*, together with the change-of-variable Jacobian, image filter, and local phase-intercept correction, gives the weighted spectrum *F*_*t,n,j*_(*p*). Different intervals have different maps, represented by the stacked spectra in Fig. 1(e); the white traces follow fixed input frequencies.

The source-to-pixel distance *L*_tx,*t*_ differs from the radial reference path 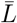 by 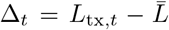. The phase-screen implementation represents this offset correction through spectral factors *A*_*s,n*_, spatial weights *B*_*s*_, and a radial query *ρ*. For one fixed event and interval, suppressing *t, j*,

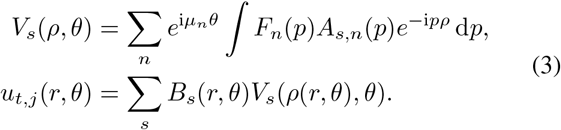

The sums and integrals use the retained terms, modes, and spectral support. The query is *ρ* = *r* + *η*(Δ_*t*_ *− ℓ*^*c*^), where the common slope *η* and path center *ℓ*^*c*^ are prepared from geometry. Each *A*_*s,n*_ weights a spectrum before synthesis; each *B*_*s*_ weights its field after evaluation at the query radius. All modes and terms share this radius for a given pixel, event, and interval [(c), (f)]. Thus the coordinate change and the weighted field sum together account for the offset phase. Panel (f) shows three contribution magnitudes from a six-term example and the magnitude of the full complex sum.

The semicircular scalar implementation samples one reference image 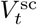 at a compensated radius [(g)]:

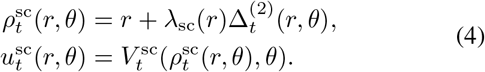

Here 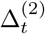 is the second-order approximation of the path difference from a centered source, and *λ*_sc_ converts that path difference into a radial displacement through a geometry-only fit. The reference image uses the scalar implementation’s radial slope and includes modal interval overlap before this query. In both implementations, the query changes where a field is read; the output pixel stays at (*r, θ*).

For phase screens, normalized interval weights 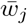 give 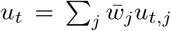. Either implementation then supplies event images for *I* = Σ_*t*_ *v*_*t*_*u*_*t*_, with normalized transmit weights *v*_*t*_. Envelope detection and logarithmic compression follow this coherent complex sum [(h)]. Panels (a)–(f) and (h) illustrate one C5-2v dataset; (g) is an independent semicircular example.

### B. Matched Inputs and Image Coordinates

Each DAS/CAMM pair uses the same RF, image grid, propagation sound speed, frequency band, and declared acquisition weights. The interface reads element coordinates, transmit delays and apodization, receive sample ranges, and sampling frequency. It splits the RF buffer into individual firings and applies the pulse, receive-start, and source-delay corrections to establish a common propagation clock. DAS linearly interpolates the conditioned analytic traces at the source-to-pixel-to-receiver travel times. CAMM incorporates the same time origin into its temporal spectra.

The comparison evaluates these complete reconstruction pipelines with their stated numerical settings. Their interpolation, modal weighting, and approximation choices remain part of the implementations being assessed.

For C5-2v, the Cartesian origin is at the probe apex, and the curvature center is (0, *− R*), where *R* is the probe radius. Native polar coordinates satisfy *x* = *r* sin *θ* and *z* = *r* cos *θ − R*; radial depth is *d* = *r − R*. Thus radial depth *d* and Cartesian depth *z* differ away from the probe axis. Semicircular coordinates are measured from the curvature center, so *x* = *r* sin *θ* and *z* = *r* cos *θ*. Measurements use image arrays before display resampling.

### C. Geometry-Dependent Precomputation

The retained frequency band and angular modes, radial intervals, interpolation maps, query coordinates, and spatial weights are prepared once for fixed acquisition and processing settings. RF-dependent spectra and fields are recomputed for each frame. Radial synthesis uses a dense matrix, a low-rank matrix product, or a nonuniform FFT (NUFFT) [21]. These backends evaluate the same prepared Stolt samples; the low-rank and NUFFT results are checked against dense evaluation. The GPU batches transmit–term pairs within each radial interval. Supplementary Section S4 details backend selection and storage.

The following section specifies the two acquisition geometries, reconstruction settings, and evaluation measures. Results then compare point-response shape, low-echo target separation, and execution cost under those fixed protocols.

## III. Simulation and Experimental Design

Table I summarizes the two acquisitions, and Table II groups the studies by simulation and measurement. The two simulated geometries serve complementary purposes. The semicircular array provides controlled sweeps of cyst position, size, and transmit count over a broad angular aperture. The convex-array simulations reproduce the C5-2v acquisition used in the experiments, linking known-target responses across depth and angle to phantom and carotid measurements. Each study compares DAS with the CAMM implementation specified for that acquisition.

**TABLE I.** Probe and Transmit Parameters.

| Parameter | Semicircular | C5-2v |
|---|---|---|
| Receive elements | 64 | 128 |
| Curvature radius (mm) | 0.75 | 49.57 |
| Virtual-source radius (mm) | 0.375 | 24.785 |
| Angular aperture (°) | 180 | 60.919 |
| Center frequency (MHz) | 20 | 3.57 |
| RF sampling rate (MHz) | 200 | 13.8889 |
| Sound speed (m/s) | 1540 | 1540 |
| Transmit mode | Diverging virtual sources |  |
| Nominal transmit count | 24 | 24 |
| CAMM offset correction | Scalar pullback | Phase screens |

**TABLE II.** Overview of Simulation and Experimental Studies.

| Study | Main question and varied parameter | Results |
|---|---|---|
| *Simulation studies* |  |  |
| Semicircular geometry | Cyst contrast versus angular position, axial position, and radius | Fig. 2 |
| Semicircular transmit count | Cyst contrast versus number of diverging transmissions | Fig. 2 |
| C5-2v points | Response width and energy distribution versus depth and angle | Fig. 3 |
| C5-2v cysts | Contrast versus depth, angle, and size; variation across scatterer realizations at fixed on-axis positions | Fig. 4, S3 |
| *Experimental studies* |  |  |
| Phantom reflectors (N1–N5) | Response width and local energy distribution versus depth and lateral position | Figs. 5, 6 |
| Phantom cysts | Inclusion–background contrast in C1/C2 and the shallow C0 region | Fig. 7; Table III; S2 |
| Carotid views | Lumen–tissue separation in longitudinal and transverse views | Fig. 8; Table IV |

### A. Semicircular Array Simulations

The semicircular study evaluates cyst contrast under changes in target geometry and transmit count. The array has 64 elements on a 0.75-mm-radius arc and virtual sources at radius 0.375 mm. Field II channel data were generated at a centre frequency of 20 MHz and sampled at 200 MHz. Both methods use the same 8–24-MHz spectrum, a 220-ns pulse-centre correction, and trapezoidal transmit weights. The receive angles are the centred element coordinates of the simulation. CAMM retains |*n*| *≤* 30 and the semicircular scalar radial pullback illustrated in Fig. 1(g). Its Lanczos modal weights are sinc(*n/*31), with sinc(*x*) = sin(*πx*)*/*(*πx*) and value one at zero. Complex radial fields are phase-deramped before interpolation and their carriers restored at the query radius.

Each phantom contains 10^5^ scatterers in the half-annulus 2 *≤ r ≤* 7 mm, with no scatterers inside the circular cyst. The baseline cyst has centre (0, 4.25) mm and radius *a*_*c*_ = 0.75 mm. Three six-value sweeps vary angular position *θ*_*c*_ *∈ {*0, 8, 16, 24, 32, 40 *}*° at fixed *z*_*c*_ = 4.25 mm, axial position *z*_*c*_ *∈ {* 3, 3.5, 4.25, 5, 5.75, 6.5*}* mm at *x*_*c*_ = 0, or radius *a*_*c*_ *∈{* 0.40, 0.60, 0.75, 0.90, 1.10, 1.30*}* mm. For the angular sweep, *x*_*c*_ = *z*_*c*_ tan *θ*_*c*_. The three sweeps share one baseline, giving 16 distinct configurations. Each configuration uses one generated phantom with the same random seed; the sweep curves therefore report one realization per configuration. Each configuration is reconstructed on the same 220 *×* 360 polar grid, covering *r* = 1.5–7.5 mm and *θ ∈* [−90°, 90°).

The cyst ROI has radius (0.55*/*0.75)*a*_*c*_; a separate back-ground circle lies within the scattering region. The same physical ROIs and equally spaced Cartesian samples are used for both methods, following Section III-D.

A separate transmit-count study measures the gain from additional illumination at the baseline geometry. It uses nine speckle realizations with 4864 scatterers each and the same baseline cyst geometry. Each realization is held fixed while the number of uniformly distributed source angles varies over *{*4, 6, 8, 10, 12, 14, 16, 20, 24 *}*. Each count spans the same angular endpoints; the source positions are redistributed uniformly for each count. The grid has 220 *×* 360 samples over *r* = 1.5–7 mm. A 0.55-mm cyst ROI is compared with four equally sized background ROIs at *r* = 4.25 mm and angles *±*30°, *±*50°. The curves report the mean and sample standard deviation over the nine realizations at each transmit count; CNR is converted to dB before aggregation.

### B. Acquisition-Matched Convex-Array Simulations

Known point and cyst targets are simulated under the C5-2v geometry and firing sequence used for the experiments. The recorded element coordinates, transmit delays, and apodization define the 128-element aperture and 24 diverging transmissions in Table I. Element width is approximately 0.391 mm, center spacing is 0.415 mm, and elevation aperture and nominal focal distance are 13.5 and 60 mm. Field II evaluates the finite rectangular-element responses [22], [23].

Both C5-2v reconstructions retain 2.15–4.99 MHz with the same frequency taper and receive-endpoint taper. CAMM retains 127 angular modes and uses exact virtual-source distances, a 0.3-rad criterion for local radial phase linearization, and a 10^−3^ phase-screen truncation tolerance, with at most 24 retained terms. These settings also apply to the recorded C5-2v data.

The recorded Verasonics excitation and impulse-response arrays specify the pulse model. Expanding the excitation states onto a 250-MHz clock and convolving with the one-way impulse response reproduces the recorded one-way waveform. Point-target RF is generated at 250 MHz, antialias-filtered, and decimated by 18 to 13.8889 MHz. Cyst simulations use the resampled pulse at 83.3333 MHz and decimation by six to the same RF rate. Against the 250-MHz reference in five point tests spanning depth and angle, this lower internal rate changes band-normalized RF by at most 0.493% in relative *L*_2_ and response widths by at most 0.156%, with unchanged peak coordinates. Each firing retains the recorded 3200-sample receive window; the reconstruction origin includes the pulse and lens delays used in synthesis.

The point study samples radial depths *d* = *{*10, 15, 20, 30, 50, 100*}* mm and angles *θ* = *{*0, −10, −20, −25 *}*°. Each of the 24 locations is a separate single-scatterer simulation with the same acquisition, pulse, and scattering amplitude. Each physical element uses 64 elevation subdivisions. Increasing this count to 128 at the 50-mm on-axis point leaves peak coordinates unchanged and changes widths by less than 0.04%.

Full-field images use the recorded-data 512 *×* 513 grid; CAMM reuses its plan across the 24 datasets. Fine-grid reconstructions are centered on the prescribed targets and use 12 *×* 8-mm tangential–radial windows at 0.025-mm spacing, with separately prepared intervals and unchanged RF, band, modes, and tolerances. At 10 mm and −25°, the local grid meets the angular boundary; this boundary-limited window is excluded from the local energy comparison in Supplementary Section S1. Figure 3 pairs representative local images with six-depth metric curves at all four angles. Its overview is a display crop; full-field fractions in (5) are calculated on the uncropped images.

Angular amplitude changes are measured by the full-field envelope maximum relative to the same method’s on-axis maximum at the same depth, expressed in dB. Supplementary Section S1 specifies this normalization and reports the position-dependent values.

Eighteen configurations use one shared 20 *×* 20-mm tangent– radial template of 24 000 uniformly randomly positioned scatterers with Gaussian amplitudes, set to zero inside a radius-*a* circle. The template is rotated and translated without redrawing scatterers. Sixteen cases use *a* = 4 mm at *d* = *{*10, 15, 30, 50*}* mm and *θ* = *{*0, − 5, −10, −20*}* °; two size controls use *a* = 2 mm at 30 mm, on axis and at −20°.

All cases use 64 elevation subdivisions, 24 transmissions, and the same 512 *×* 513 image grid. The internal ROI radius is 0.8*a* and the background annulus spans 1.3*a*–1.8*a*. Metrics use uncompressed envelopes, polar pixel-area weights, and 64 shared histogram bins after each method’s background-mean normalization. Figure 4 shows paired images at −20° and metric differences at all sixteen radius-4-mm locations.

An on-axis cyst study measures variation across scatterer realizations at fixed positions. Radius-4-mm cysts at (*x, z*) = (0, 25), (0, 60), (0, 100) mm are evaluated with five scatterer realizations per depth, using the same acquisition and ROI protocol. Supplementary Section S3 reports metrics for the 18 spatial configurations and the on-axis means and sample standard deviations.

### C. Phantom and In Vivo Experiments

The recorded-data study tests whether the spatial-response and contrast trends recur in phantom and carotid images. Image-quality evaluation uses eight RF records acquired with a Verasonics Vantage platform and C5-2v probe using 128 receive channels: five cross-target phantom records N1–N5 at different lateral placements, one phantom record containing cysts C1 and C2, and two carotid views from one of the authors. Each record contains 20 buffer frames with 24 diverging transmissions per frame. Each DAS/CAMM pair uses the same RF, recorded geometry, firing sequence, band, and declared acquisition weights. Buffer frame 1 supplies the representative images. Medians and minimum–maximum ranges summarize variation within each record.

The KS107BG(N-3) small-organ tissue-mimicking phantom was manufactured by the Institute of Acoustics, Chinese Academy of Sciences. Its label specifies a sound speed of 1540 *±* 10 m/s, attenuation of 0.70 *±* 0.05 dB/(cm MHz), and reference temperature of 23 *±* 3 °C. RF was saved for offline reconstruction on the workstation specified in Section IV-G; the acquisition console and reconstruction workstation are separate systems.

### D. Image-Quality Evaluation

Point and cyst measures describe complementary spatial scales. Speckle and contrast-detail studies relate visibility to target size and background fluctuations [24], [25], while cystic-resolution measures connect spatial response to contrast [26], [27]. Metrics use the envelope *A* = |*I*| before log compression [28]. Radial and tangential FWHM use linearly interpolated half-amplitude crossings on fine-grid reconstructions. Tangential width is *r*Δ*θ*, using the prescribed target radius in simulations and the detected peak radius in measurements. Simulated localization error is measured from the prescribed target to the peak in its search neighborhood; measured DAS– CAMM peak distance describes disagreement between the two reconstructions.

The outside-neighborhood energy fraction measures the distribution of squared image magnitude beyond a fixed neigh-borhood *N*:

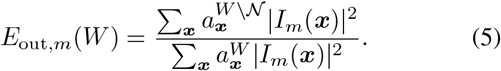

Here *m* denotes the method and *a*^*D*^ the polar pixel-area weight assigned to domain *D*. The inner neighborhood spans *±*1 mm radially and *±*3 mm tangentially about the reference target. Full-field *E*_out_ uses the entire reconstruction as *W* and pixel-center masks. Local *E*_out_ uses a 12 *×* 8-mm tangential–radial window and fractional boundary-cell areas. Both use full-field complex images and identical physical masks within a pair.

FWHM measures the central peak, whereas *E*_out_ describes energy distribution over a larger domain: a peak can broaden within *N* while the fraction outside it decreases. Both numerator and denominator depend on the reconstructed image, so the fraction expresses exterior energy relative to the total energy in *W*. Measured windows also contain background echoes. Full-field and local values are reported separately because their integration domains differ.

For cyst and background ROIs, *µ*_*c*_, *µ*_*b*_ are envelope means, 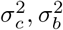 are pixel-distribution variances, and *p*_*c*_, *p*_*b*_ are normalized envelope distributions. The contrast measures are

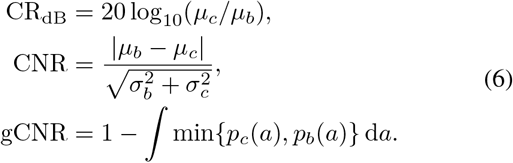

For low-echo targets, more negative CR, higher CNR, and higher gCNR indicate improvement. CR describes the mean envelope ratio; CNR additionally accounts for within-region fluctuations; gCNR describes distributional separation [29]. Their rankings can therefore differ. CNR is dimension-less: simulation plots labeled CNR (dB) use 20 log_10_(CNR), whereas measured-data tables report linear CNR. The gCNR histogram estimate depends on sample support and bin selection [30].

Semicircular ROI samples lie on an equal-area Cartesian grid; background samples from multiple ROIs are pooled to calculate population moments. Convex studies use polar pixel-area-weighted means and variances. Both methods use the same physical masks. Each envelope is divided by its own background mean before forming paired 64-bin histograms with shared edges. Display normalization is specified separately in each figure.

For recorded reflectors, scanner-reference images locate fixed target windows, and independent fine-grid RF reconstructions supply widths on paired valid frames. Local energy fractions use all frames in complete windows without neighboring targets. C1/C2 regions are fixed from the scanner-reference image, and carotid masks sample the lumen interior and adjacent tissue. All masks remain fixed across methods and stored frames. Supplementary Section S2 specifies window eligibility, the RF limit-sample criterion, fixed ROI coordinates, and integration and histogram settings.

### E. Computational Measurements

Timings use an Intel Core i7-13700KF workstation with 64 GiB RAM and an NVIDIA RTX 4090 (24 GiB), running Windows 11 Pro, Python 3.11.4, and CuPy 13.6.0. Host latency runs from an RF frame in host memory to the completed complex image returned to host memory. It includes transfers and reconstruction, excluding file loading, B-mode rendering, and one-time plan preparation. Medians summarize 16 executions per method after warm-up; the protocol is given in Supplementary Section S4.

## IV. Results

### A. Semicircular Cysts

#### 1) Cyst position and radius

Figure 2(a) shows the baseline geometry, and (b)–(d) illustrate the three geometric sweeps; the highlighted configurations correspond to the image pairs below them. Black and orange arcs denote the receive and virtual-source loci, and the dotted arc marks the outer scattering boundary. The images use full-field peak normalization over a 60-dB range; dashed circles mark the evaluation ROIs.

**Fig. 2.**
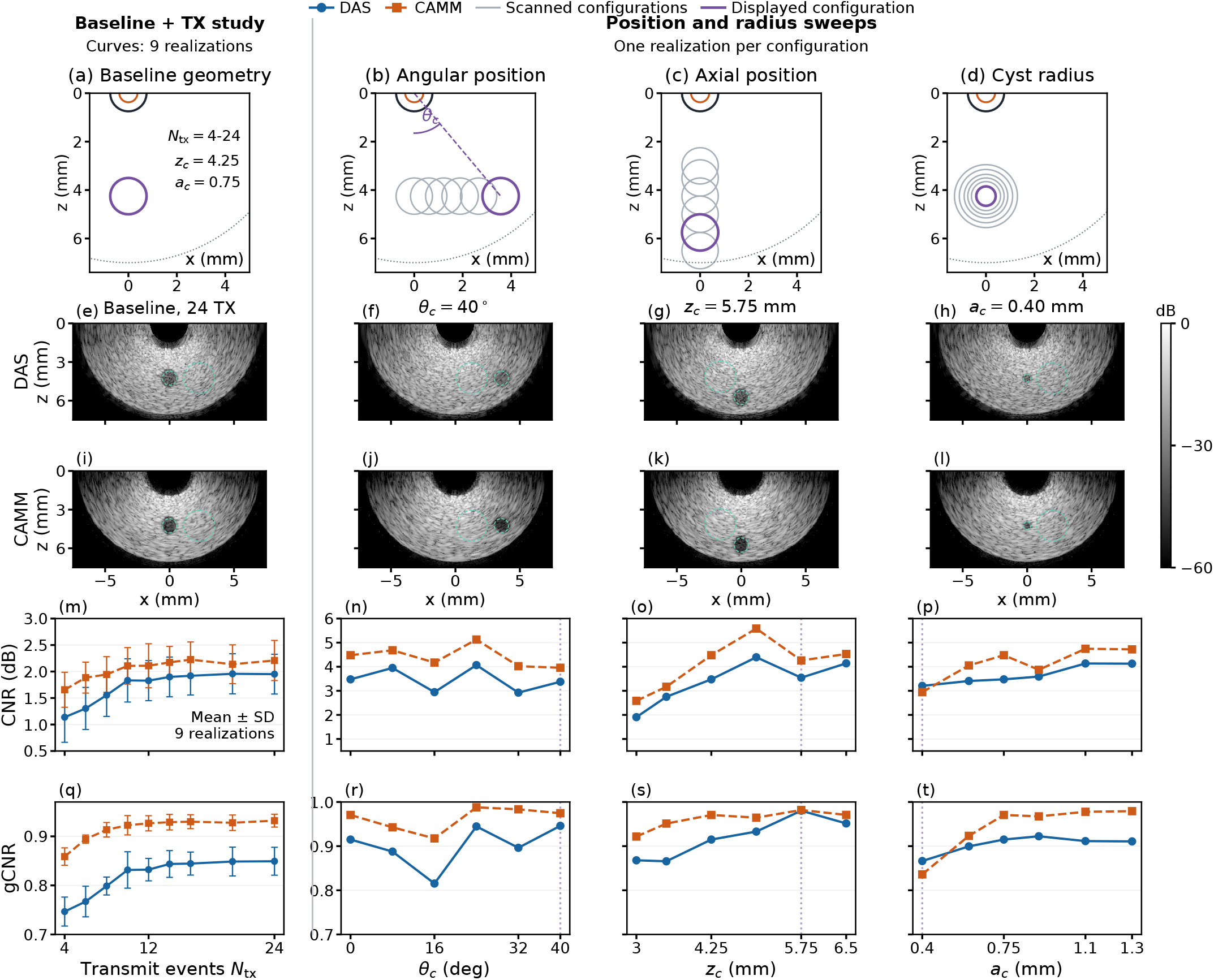
Semicircular cyst studies. (a)–(d) Geometry; purple marks the displayed configurations and gray the scanned configurations; (b) varies *θ*_*c*_ at fixed *z*_*c*_. (e)–(l) DAS and CAMM images at 24 transmissions. (m)–(p) CNR (dB); (q)–(t) gCNR. First-column curves: mean *±* sample SD over nine scatterer realizations, with independent vertical scales. Columns 2–4: angular-position, axial-position and radius sweeps, with one scatterer realization per configuration. Images use each full-field peak as the 60-dB display reference.

**Fig. 3.**
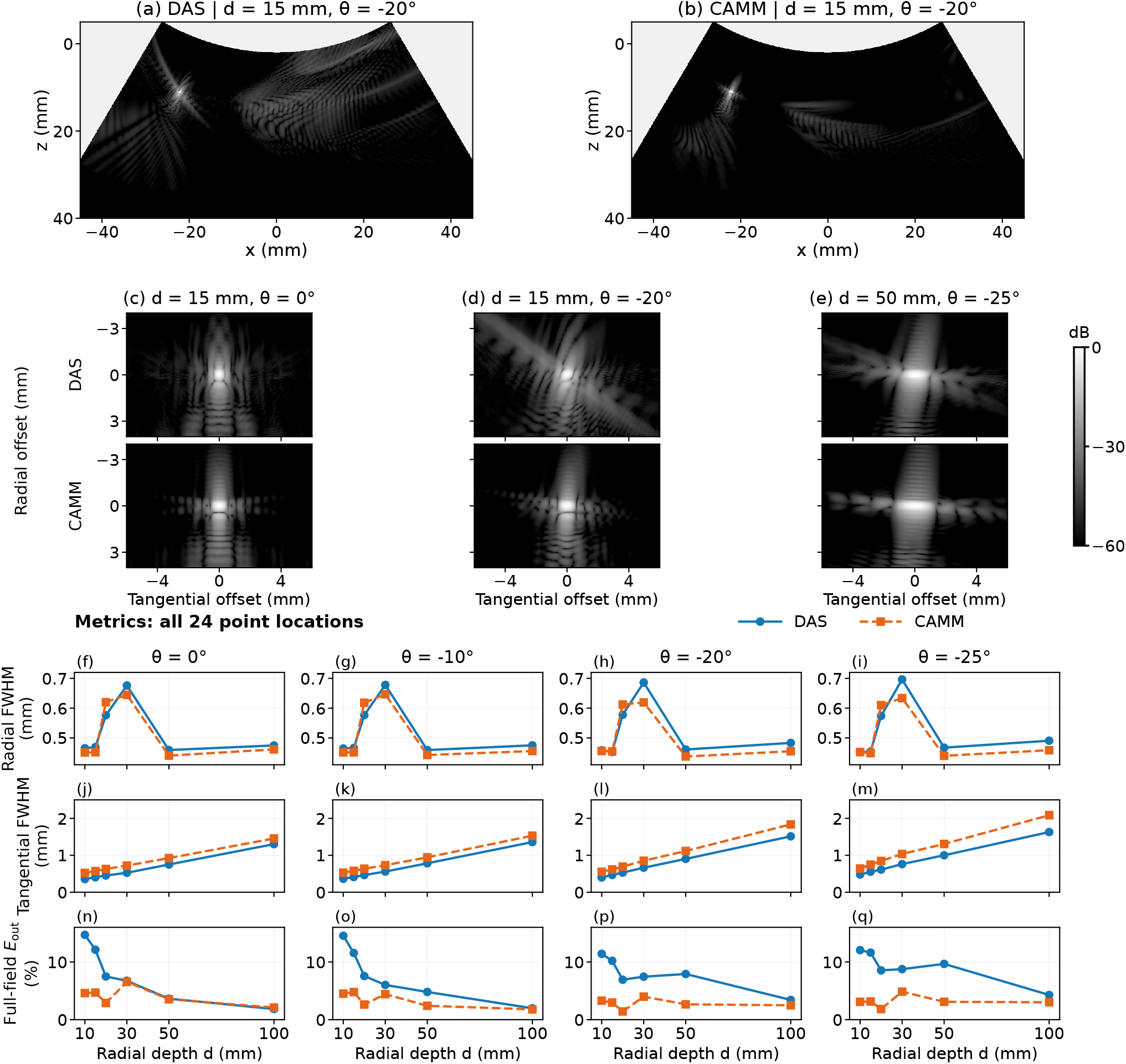
C5-2v single-point responses. (a,b) Overview at *d* = 15 mm, *θ* = −20°. (c)–(e) Paired local images at (*d, θ*) = (15 mm, 0°), (15 mm, −20°), and (50 mm, −25°). (f)–(i) Radial FWHM; (j)–(m) tangential FWHM; (n)–(q) full-field *E*_out_ at all six depths and four angles. Local windows span 12 *×* 8 mm; each image uses its own reconstruction peak over 60 dB. Energy fractions use the uncropped full field.

The baseline pair in Fig. 2(e), (i) and the position examples in (f), (j) and (g), (k) show lower cyst-interior echo levels in CAMM relative to the surrounding background. The corresponding curves in (n)–(p), (r)–(t) show higher CAMM CNR and gCNR in 15 of the 16 distinct configurations. At the baseline, CNR increases from 3.476 to 4.478 dB and gCNR from 0.915 to 0.971. The smallest cyst, *a*_*c*_ = 0.40 mm, shown in (h), (l), reverses that ranking: CNR changes from 3.205 to 2.945 dB and gCNR from 0.866 to 0.836 for DAS and CAMM, respectively. The contrast benefit therefore depends on the target scale as well as its position.

#### 2) Number of transmissions

Figure 2(m), (q) shows increasing mean contrast metrics followed by smaller changes at higher transmit counts. From 4 to 24 transmissions, mean gCNR rises from 0.747 to 0.849 for DAS and from 0.859 to 0.932 for CAMM. CAMM reaches 0.927 at 12 transmissions; its mean CNR changes from 2.110 to 2.209 dB between 12 and 24, compared with 1.828 to 1.951 dB for DAS. For this fixed-endpoint acquisition plan, the main improvement occurs before 12 transmissions. Each count uses a newly uniform distribution of source angles, so these comparisons describe a set of acquisition plans with common angular endpoints.

The absolute CNR levels are lower than in the geometric sweeps, which use a different background realization and sampling scheme. This ensemble has about 21 times fewer scatterers and samples the background through four small ROIs; the geometric sweeps use one larger background ROI. For CAMM at 24 transmissions, the background standard-deviation-to-mean ratio averages 0.717, versus 0.551 for the geometric baseline, while the cyst-to-background mean ratio is approximately 0.075 in both. Greater relative background variation increases the CNR denominator. This observed back-ground variation accompanies changes in both scatterer density and ROI layout. The independent vertical scales in Fig. 2(m), (q) emphasize the within-study transmit-count trend.

### B. C5-2v Point Responses

CAMM has a lower full-field *E*_out_ at 23 of 24 positions, with the largest reductions at the sampled shallow depths. At the displayed 15-mm, − 20° point, the fraction decreases from 10.20% for DAS to 2.97% for CAMM, alongside visibly weaker extended curved responses. Across the 12 locations at 10–20 mm, the decrease is 4.59–10.09 percentage points. The exception is 100 mm on axis, where the fraction increases from 1.85% to 2.10%.

The central peak shows a different tradeoff. CAMM has narrower radial FWHM at 18 of 24 positions but broader tangential FWHM at all 24. At 50 mm on axis, radial width changes from 0.459 to 0.441 mm, while tangential width increases from 0.750 to 0.924 mm. All four 20-mm targets have broader CAMM radial peaks despite lower full-field exterior fractions. The curves therefore distinguish changes in the central peak from changes in the fraction of power outside the fixed neighborhood.

Using the local 12 *×* 8-mm domain in (5), CAMM has a lower local *E*_out_ at 20 of the 23 complete windows; the 10-mm, −25° window is excluded because it meets the field boundary. Supplementary Section S1 provides the same-window comparison. Fine-grid localization errors reach 0.071 mm for DAS and 0.056 mm for CAMM; on the full-field grid, the maximum is 0.115 mm for both.

Both methods’ on-axis-normalized peaks decline toward the angular edge. At − 25°, CAMM’s relative peak is slightly higher at 10 mm and lower at the sampled 20–100-mm depths; Supplementary Section S1 reports the corresponding normalized peak values.

### C. C5-2v Cyst Contrast

The point study shows lower exterior fractions alongside broader tangential peaks. We next test whether low-echo targets are better separated from a speckled background.

The shared-realization spatial scan shows a consistent shallow-region benefit and mixed deeper results. All eight radius-4-mm configurations at 10–15 mm favor CAMM in CR, CNR, and gCNR. Among the eight configurations at 30– 50 mm, two favor CAMM in all three measures, two favor DAS, and four give mixed rankings. The difference maps in Fig. 4 summarize these spatial changes; their colors indicate the direction of each paired difference.

**Fig. 4.**
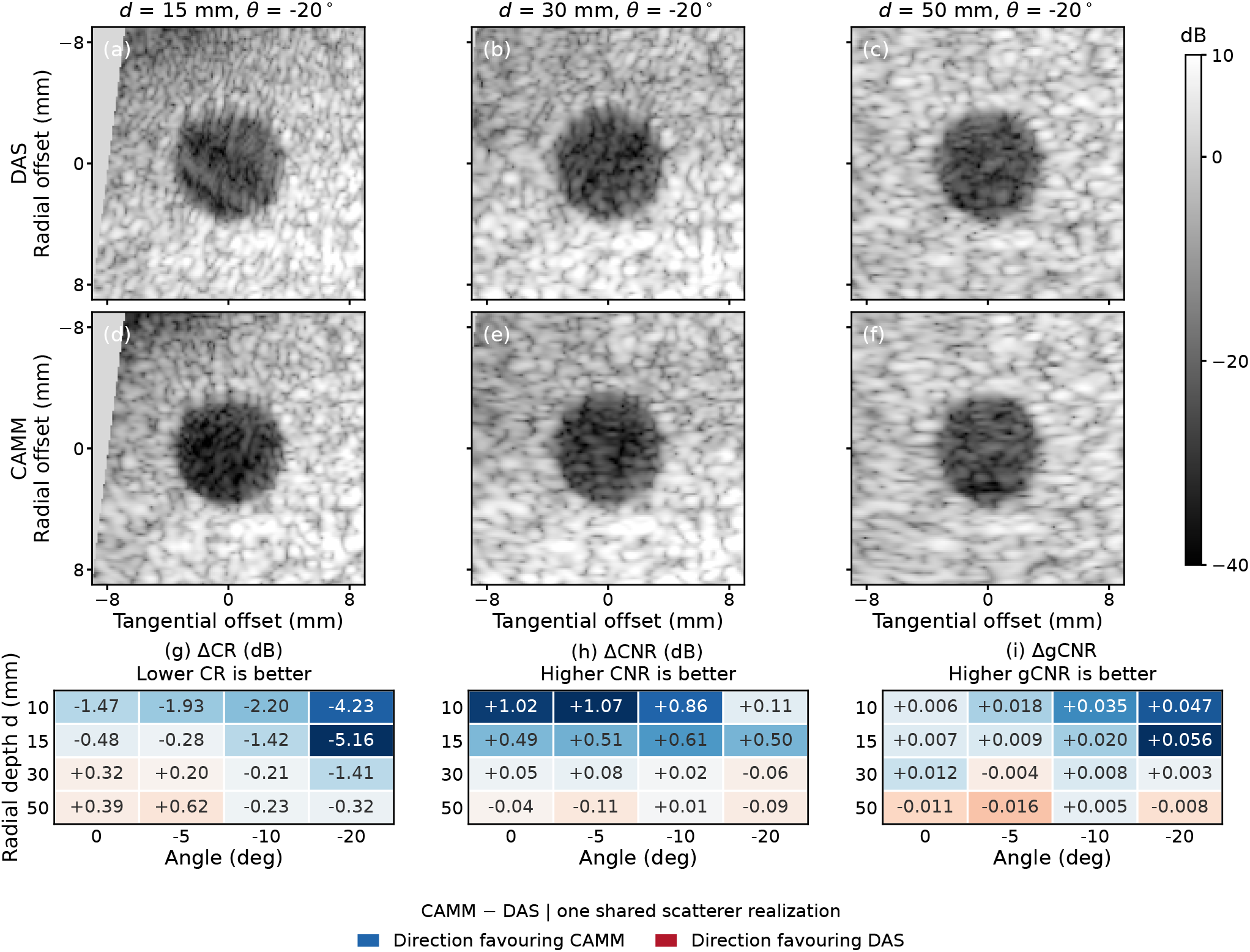
Radial-depth–angle cyst comparison. (a)–(f) DAS above CAMM at −20° in 18 *×* 18-mm windows, displayed from − 40 to 10 dB relative to each method’s background mean. (g)–(i) CAMM-minus-DAS differences at sixteen radius-4-mm positions; blue favors CAMM, red DAS. Lower CR and higher CNR/gCNR are favorable; CNR differences are in dB. These configurations share one scatterer realization.

In the 15-mm, −20° image pair, CAMM has a darker inclusion against the surrounding speckle: CR changes from − 22.64 to −27.80 dB, CNR from 1.40 to 1.90 dB, and gCNR from 0.901 to 0.957. The deeper cases illustrate why the readouts are complementary. At 30 mm and −5°, CNR increases by 0.08 dB, while CR becomes 0.20 dB less negative and gCNR falls by 0.004. At 50 mm, all three measures favor DAS on axis and at −5°, whereas the off-axis comparisons are mixed.

The radius-2-mm controls also separate the readouts. At 30 mm and −20°, CR becomes 2.25 dB more negative and gCNR rises by 0.016, but CNR decreases by 1.02 dB. On axis, CR and CNR favor DAS while gCNR favors CAMM. Thus the spatial scan identifies a shallow-region contrast benefit for this shared realization, with deeper and smaller-target results depending on the selected metric.

The on-axis study in Supplementary Section S3 quantifies contrast variation over five scatterer realizations at each depth. Mean gCNR is 0.955/0.949, 0.928/0.922, and 0.944/0.940 for DAS/CAMM at 25, 60, and 100 mm, respectively. Mean CNR increases with CAMM at 25 mm and decreases slightly at 60 and 100 mm.

### D. Measured Phantom Responses

The schematic in Fig. 5 identifies the evaluated target groups. The paired images show the cross-target layout in N5 and N3, placing the vertical target column near the axis and to its right, respectively. The two horizontal rows provide lateral samples near 19 and 39 mm. The cyan window locates the shallow candidate inclusion region C0. Each image uses its full-field peak as the display reference. Supplementary Section S2 summarizes the measurement protocol and record-level reflector results for N1–N5.

**Fig. 5.**
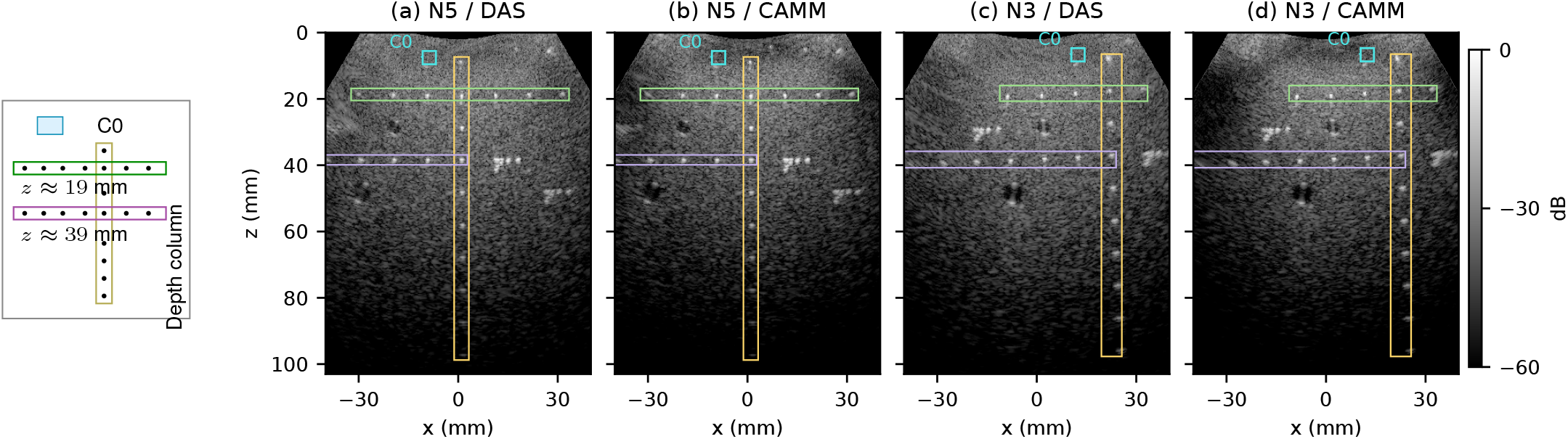
Cross-target phantom. Left: schematic of evaluated target groups. Images: N5 (a,b) and N3 (c,d), buffer frame 1. Yellow: vertical column; green and purple: horizontal rows; cyan: C0 region. The N5 rows provide the lateral samples in Fig. 6(b,d,f); the depth curves also include other records. Each DAS/CAMM image uses a 60-dB scale relative to its own full-field peak.

#### 1) Depth- and position-dependent reflectors

Across the five records, larger CAMM tangential widths coexist with lower local energy fractions. Median tangential FWHM is larger in 91 of 94 isolated target windows, while median local *E*_out_ is lower in all 92 eligible energy windows. The larger-window neighbor-exclusion rule removes two windows from energy evaluation. RF limit-sample flags identify 53 target echo windows, all retained in the eligible comparisons. The same trends hold in all 41 unflagged width windows and all 39 unflagged energy windows; Supplementary Section S2 gives the flag and eligibility rules.

Figure 6(a,c) shows stronger depth dependence in the tangential response than in the radial width. Along the unflagged N5 column at *z* = 48.1–97.3 mm, radial widths remain about 0.54–0.60 mm; tangential widths rise from 0.912 to 1.752 mm for DAS and from 1.028 to 1.790 mm for CAMM. The common width rule retains 16–20 paired frames per window; Supplementary Section S2 specifies the paired-frame validity rule.

**Fig. 6.**
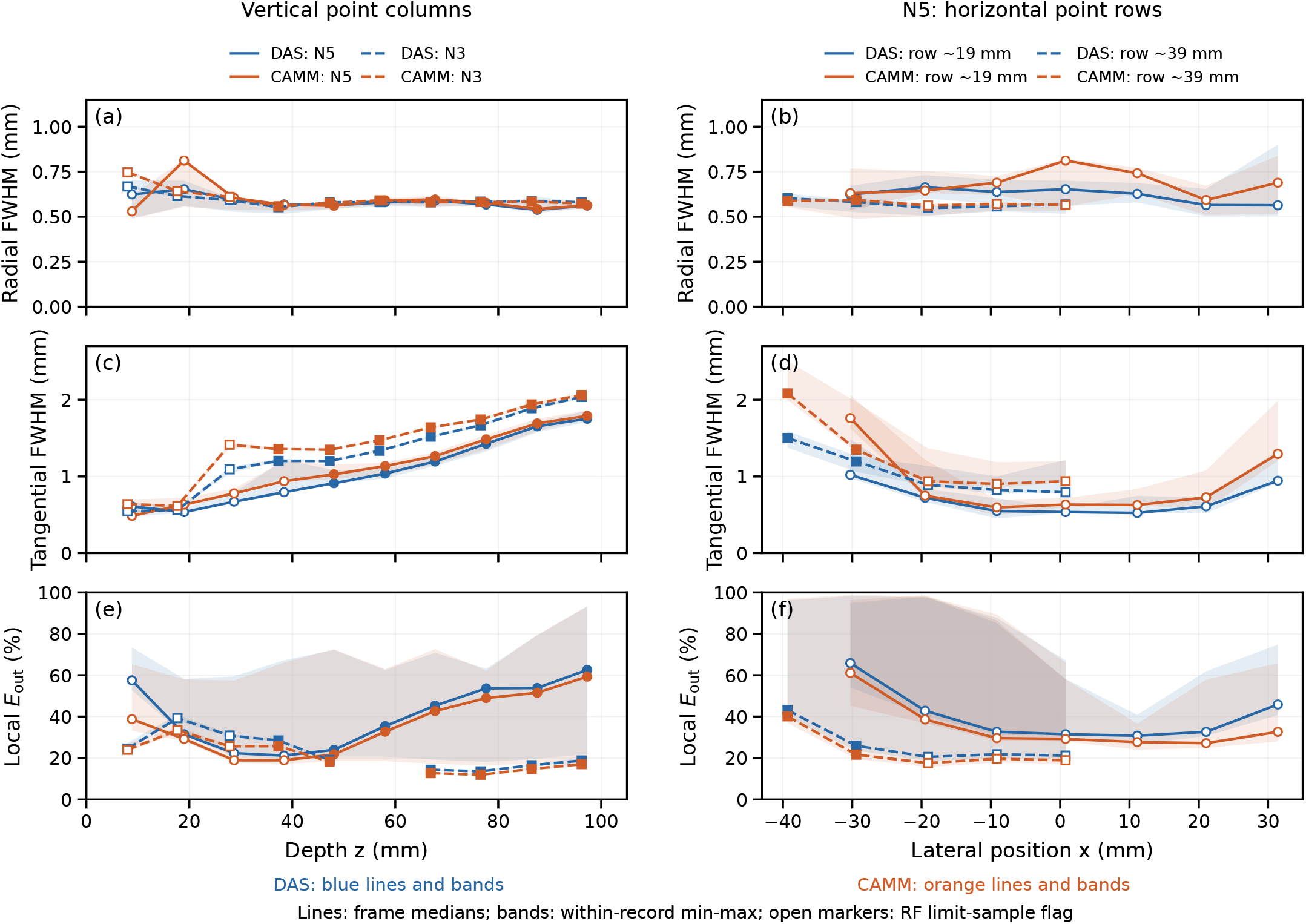
Measured reflector responses versus Cartesian depth *z* (left) and lateral position *x* along the two N5 rows (right): radial/tangential FWHM and local *E*_out_. Blue denotes DAS and orange denotes CAMM. Lines give within-record medians; matching shaded bands span framewise minima– maxima over paired valid frames for widths and all 20 frames for fractions. Open symbols identify predicted target echo windows containing RF samples at the recorded amplitude limits; gaps mark excluded energy windows.

The N5 horizontal rows in panels (b,d) show narrower responses near the center and unequal broadening toward the two sides. At the unflagged leftmost point of the approximately 39-mm row, tangential FWHM is 1.502 mm for DAS and 2.083 mm for CAMM, while radial widths remain about 0.58– 0.60 mm. At the far-left N3 target near (−38.5, 39.3) mm, the asymmetric CAMM response also changes the selected peak: the median DAS–CAMM peak distance is 0.665 mm and its maximum is 1.252 mm. This distance measures disagreement between the two reconstructions, whereas the simulated localization errors use the known target position.

Panels (e,f) quantify the local energy-fraction changes. For each window, we first take the median of its 20 framewise CAMM-minus-DAS differences. The median of these window summaries is −2.85 percentage points overall and −2.59 in the unflagged subset. Thus the measured response changes involve both a broader central peak and a lower exterior fraction, with additional asymmetry and peak disagreement at some peripheral targets.

### E. Measured Phantom Cysts

The response measurements characterize local reflector shapes; we now examine separation of low-echo inclusions from background. Both C1 and C2 show more negative CR and higher CNR and gCNR with CAMM in all 20 paired frames (Table III). The magnitude differs between the two regions.

**TABLE III.** Measured Cyst Contrast: Within-Record Medians (20 Frames)

| Target | Method | CR (dB) | CNR (linear) | gCNR |
|---|---|---|---|---|
| C1 | DAS | -8.68 | 1.140 | 0.681 |
|  | CAMM | -10.11 | 1.300 | 0.754 |
| C2 | DAS | -4.73 | 0.790 | 0.541 |
|  | CAMM | -17.27 | 1.616 | 0.900 |

For C1, Fig. 7(a,b) shows similar surrounding speckle and a modest reduction in inclusion echo. Median CR changes from −8.68 to −10.11 dB and gCNR from 0.681 to 0.754. C2 has a more pronounced change: residual echo remains inside the DAS inclusion, whereas CAMM gives a darker interior against adjacent tissue. Median CR changes from −4.73 to −17.27 dB and gCNR from 0.541 to 0.900, corresponding to histogram overlap decreasing from 0.459 to 0.100. C2 gCNR ranges are 0.508–0.580 for DAS and 0.878–0.915 for CAMM, showing that the representative-image difference persists within the acquisition.

**Fig. 7.**
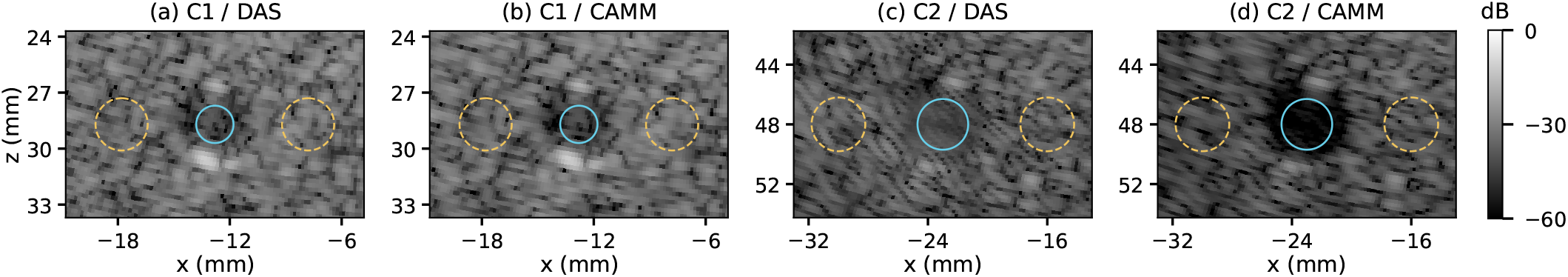
Measured cysts: (a,b) C1 and (c,d) C2, DAS/CAMM, buffer frame 1. Cyan: interior ROIs; dashed yellow: tissue references. Images use a 60-dB scale relative to each full-field peak.

The layout-indicated C0 region was also reconstructed on a 25-*µ*m grid. Its indistinct boundary makes the five fixed-window measurements exploratory (Supplementary Section S2). The nominal N3 window gives median CNR of 0.431 for DAS and 0.831 for CAMM, and gCNR of 0.275 and 0.520. Across the tested ROI radii and center shifts, and the additional gCNR bin-count checks, metric rankings reverse in four of the five records. The small-region contrast ranking therefore varies with the evaluated record and ROI settings.

### F. In-Vivo Images and Paired Measurements

Figure 8 compares longitudinal and transverse carotid views from the same participant. The deeper lumen marked by the cyan polygon in panels (a,b) corresponds to the circular vessel in panels (c,d), as identified by the acquiring author. All four panels retain the same field of view and physical scale. The evaluation samples the lumen interior and adjacent tissue in each view.

**Fig. 8.**
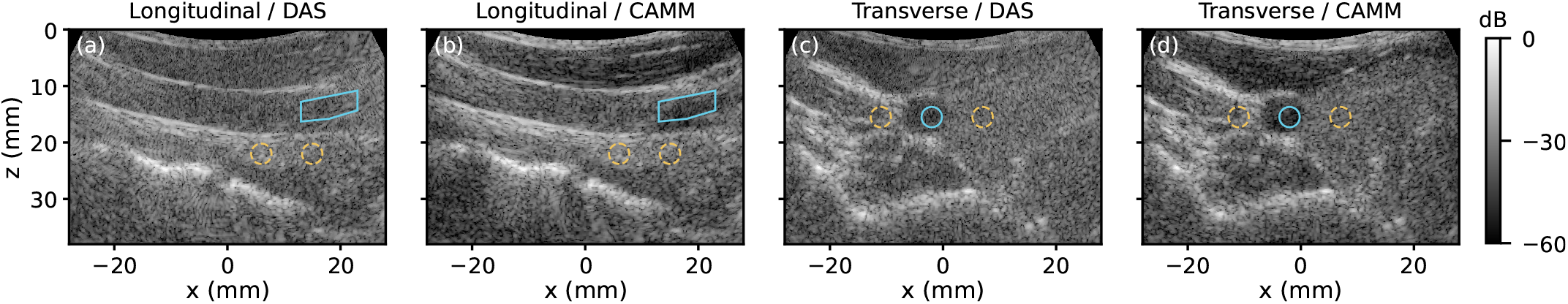
Carotid views: (a,b) longitudinal and (c,d) transverse, DAS/CAMM, buffer frame 1. Cyan: corresponding lumen interiors; dashed yellow: tissue references. All panels share the field of view and *±*27° sector; each uses its own full-field peak over 60 dB.

Both longitudinal reconstructions delineate the proximal and distal walls, while CAMM shows lower internal echo in the marked rightward segment. With the same tissue references, median CR changes from −6.41 to −11.06 dB, linear CNR from 0.683 to 1.005, and gCNR from 0.404 to 0.622 (Table IV). The gCNR ranges are 0.269–0.476 for DAS and 0.487–0.700 for CAMM.

**TABLE 4.** Carotid Contrast: Within-Record Medians (20 Frames)

| View | Method | CR (dB) | CNR (linear) | gCNR |
|---|---|---|---|---|
| Longitudinal | DAS | -6.41 | 0.683 | 0.404 |
|  | CAMM | -11.06 | 1.005 | 0.622 |
| Transverse | DAS | -8.80 | 0.930 | 0.578 |
|  | CAMM | -13.14 | 1.131 | 0.733 |

Values are within-record medians; each view uses the same physical masks for both methods across 20 frames.

The transverse view likewise shows lower CAMM internal echo while retaining the surrounding bright interfaces. Median CR decreases from −8.80 to − 13.14 dB, linear CNR increases from 0.930 to 1.131, and gCNR from 0.578 to 0.733. The gCNR ranges are 0.521–0.621 for DAS and 0.698– 0.792 for CAMM. In both evaluated lumen regions, all three paired metrics favor CAMM in each of the 20 stored frames. The paired measurements therefore support sustained contrast gains within these two recordings of the participant.

### G. Runtime and Memory

At 512 *×* 513 pixels and 24 transmissions, median latency is 29.79 ms for DAS and 51.87 ms for CAMM. These measurements include host–device transfers and reconstruction, excluding one-time preparation, file loading, and B-mode display processing. Persistent device arrays occupy 0.26 GiB for DAS and 2.05 GiB for CAMM, including reusable work buffers. DAS is faster and uses less persistent storage at all three tested grids. Supplementary Section S4 reports the grid-dependent timings and peak allocator usage.

Within the C5-2v phase-screen implementation, low-rank evaluation reduces latency from the dense implementation’s 85.30 ms by 39.19%; all 23 radial intervals use this evaluation in the reported configuration. The accelerated and dense phase-screen implementations yield closely matching complex images, with a relative *L*_2_ difference of 5.38 *×* 10^−7^ evaluated at their original gain and phase. Geometry-dependent preparation is reused across frames with fixed acquisition and processing settings; the reported latency includes the data-dependent field synthesis for each frame.

## V. Discussion

Across the point simulations and measured reflector windows, CAMM redistributes the reconstructed response: lower outside-neighborhood energy fractions often accompany broader tangential peaks. Part I relates this behavior to the radial propagation phase assigned to sampled angular components, which can weaken coherent reinforcement away from the target [15]. The present comparisons measure the combined effect of that propagation treatment, weighting, and numerical evaluation under matched acquisition inputs. Full-field point fractions describe isolated responses over the reconstruction domain; local measured fractions also include background echoes. Peripheral measured targets further show lower relative peaks together with asymmetric or displaced responses. The agreement between accelerated and dense phase-screen images reported in Supplementary Section S4 supports evaluating these changes as properties of the implemented reconstruction.

The cyst and lumen studies connect this response redistribution to regional target separation. Scalar-pullback CAMM improves CNR and gCNR in most semicircular configurations, with the smallest cyst favoring DAS. Phase-screen CAMM gives consistent gains in the sampled shallow C5-2v spatial configurations, while deeper and smaller targets yield mixed metric rankings. The five-realization on-axis study additionally gives slightly lower mean CAMM gCNR at all three depths. Measured C1/C2 cysts and the evaluated carotid lumen regions show sustained contrast gains within their records. The larger benefit in deep C2 accompanies differences in target size, background, and illumination from the simulated cysts; these factors jointly define the comparison. The indistinct C0 region retains ROI-sensitive rankings. Together, these observations support acquisition- and target-dependent benefits. The carotid evidence covers two views of one participant, with reproducibility across probe placements and participants remaining a direction for further evaluation.

The offset realization and its computational cost are part of this application-specific choice. Part I’s error analysis and matched point comparisons support scalar pullback for the compact semicircular configuration and motivate phase screens for the evaluated convex studies. Its supplementary guidance extends the selection to other bands, source offsets, and image regions. At 512 *×* 513 pixels, low-rank evaluation reduces the C5-2v phase-screen latency from 85.30 to 51.87 ms while preserving close agreement with dense CAMM; DAS takes 29.79 ms and uses less persistent device storage. Reusable plans amortize preparation across frames, while field synthesis and its buffers determine the continuing computational load. Regional target separation, response width, and available computation therefore provide complementary criteria for selecting the reconstruction.

## IV. Conclusion

CAMM was implemented as a reusable modal reconstruction pipeline and evaluated against DAS with controlled semicircular studies, acquisition-matched C5-2v simulations, and recorded phantom and carotid data. The comparisons establish a regional image-quality tradeoff: lower outside-neighborhood energy fractions and improved low-echo target separation can accompany broader tangential responses. The shallow sampled convex configurations, the two well-defined measured cyst windows, and the evaluated carotid lumen regions showed contrast gains. Deeper simulated configurations and smaller inclusions gave size-, location-, or metric-dependent rankings. These findings connect Part I’s modal construction to spatial-response and contrast changes under the tested settings. Low-rank radial evaluation reduces the cost of dense phase-screen CAMM while preserving numerical agreement, although DAS remains faster and uses less device storage in the tested C5-2v implementation. Evaluating contrast, response width, and computational cost together therefore provides a practical basis for choosing a reconstruction for the intended imaging region.

## Data and Code Availability

Selected processed data and analysis scripts supporting the reported results may be made available by the corresponding authors upon reasonable request. The scope of access and terms of use will be agreed on a case-by-case basis.

