## Supplementary Material for "Circular-Arc Modal Migration for Ultrasound Imaging—Part II: Simulation and Experimental Evaluation in Compact Semicircular and Convex Arrays"

Zihan Huang, Zhoufeng Liu, Chuan Chen*, Haixia Qi*, and Dawei Wu*

*Corresponding authors.

#### S1. C5-2v point-response comparisons

Twenty-four single-target datasets sample six radial depths $d = r - R$ and four angles about the curvature center. Each pair uses the same RF and 24 transmissions. Table S1 reports radial and tangential half-amplitude widths on a 0.025-mm local grid, full-field and local outside-neighborhood energy fractions, and relative angular peak amplitude. The latter is $G_m(d, \theta) = 20 \log_{10}[A_m(d, \theta)/A_m(d, 0)]$, where $A_m$ is the full-field envelope maximum. Each method uses its own same-depth on-axis reference.

**Table S1:** All 24 point locations. Every paired entry is DAS/CAMM. Widths are in mm, energy fractions in %, and $G$ in dB.

| $d$ (mm) | $\theta$ (°) | Radial width | Tangential width | Full $E_{\text{out}}$ | Local $E_{\text{out}}$ | $G$ |
|---|---|---|---|---|---|---|
| 10 | 0 | 0.465/0.451 | 0.355/0.523 | 14.70/4.61 | 2.03/2.62 | 0.00/0.00 |
| 10 | -10 | 0.465/0.452 | 0.363/0.530 | 14.54/4.51 | 2.25/2.58 | -1.84/-1.74 |
| 10 | -20 | 0.459/0.456 | 0.400/0.554 | 11.40/3.29 | 3.29/2.54 | -8.87/-8.19 |
| 10 | -25 | 0.452/0.453 | 0.479/0.642 | 12.05/3.09 | – | -16.86/-16.33 |
| 15 | 0 | 0.469/0.452 | 0.405/0.572 | 12.12/4.69 | 1.69/1.74 | 0.00/0.00 |
| 15 | -10 | 0.466/0.452 | 0.411/0.579 | 11.55/4.78 | 1.93/1.85 | -1.27/-1.26 |
| 15 | -20 | 0.453/0.455 | 0.467/0.616 | 10.20/2.97 | 2.79/1.53 | -6.43/-5.85 |
| 15 | -25 | 0.454/0.449 | 0.554/0.747 | 11.62/3.15 | 4.19/1.55 | -11.79/-11.81 |
| 20 | 0 | 0.576/0.620 | 0.454/0.630 | 7.48/2.89 | 0.60/0.53 | 0.00/0.00 |
| 20 | -10 | 0.577/0.618 | 0.465/0.637 | 7.56/2.63 | 0.67/0.50 | -1.12/-1.02 |
| 20 | -20 | 0.578/0.612 | 0.531/0.693 | 6.92/1.43 | 0.99/0.26 | -5.37/-5.00 |
| 20 | -25 | 0.574/0.610 | 0.612/0.847 | 8.54/1.84 | 1.67/0.23 | -9.63/-10.17 |
| 30 | 0 | 0.676/0.645 | 0.526/0.721 | 6.73/6.55 | 2.23/1.95 | 0.00/0.00 |
| 30 | -10 | 0.678/0.647 | 0.559/0.731 | 6.02/4.43 | 2.26/1.81 | -1.13/-0.77 |
| 30 | -20 | 0.686/0.619 | 0.664/0.852 | 7.45/4.00 | 2.56/1.85 | -4.68/-4.66 |
| 30 | -25 | 0.696/0.634 | 0.762/1.035 | 8.76/4.85 | 2.85/1.91 | -7.68/-8.70 |
| 50 | 0 | 0.459/0.441 | 0.750/0.924 | 3.64/3.54 | 0.63/0.44 | 0.00/0.00 |
| 50 | -10 | 0.459/0.443 | 0.783/0.944 | 4.80/2.41 | 0.65/0.47 | -0.96/-0.74 |
| 50 | -20 | 0.461/0.438 | 0.902/1.115 | 7.91/2.66 | 0.76/0.55 | -4.26/-4.34 |
| 50 | -25 | 0.467/0.440 | 1.000/1.306 | 9.66/3.10 | 0.79/0.51 | -7.00/-7.82 |
| 100 | 0 | 0.475/0.462 | 1.302/1.455 | 1.85/2.10 | 0.76/0.52 | 0.00/0.00 |
| 100 | -10 | 0.475/0.456 | 1.357/1.530 | 1.98/1.79 | 0.78/0.57 | -0.85/-0.96 |
| 100 | -20 | 0.483/0.455 | 1.513/1.836 | 3.42/2.50 | 0.91/0.73 | -3.45/-4.37 |
| 100 | -25 | 0.491/0.459 | 1.629/2.085 | 4.28/3.02 | 0.99/0.89 | -5.47/-7.07 |

Both energy measures use the uncompressed full-field complex image and an inner neighborhood of ±1 mm radially and ±3 mm tangentially. Full-field $E_{\text{out}}$ uses all reconstructed pixels with pixel-center masks. Local $E_{\text{out}}$ uses the $12 \times 8$-mm tangential–radial window and fractional polar-cell areas specified in S2. The dash marks an incomplete local window. Full-field localization errors reach 0.115 mm for both methods; fine-grid errors reach 0.071 mm for DAS and 0.056 mm for CAMM.

CAMM has lower full-field $E_{\text{out}}$ at 23/24 locations, except 100 mm on axis. Local $E_{\text{out}}$ is lower at 20/23 eligible locations; exceptions are 10 mm at 0° and −10°, and 15 mm at 0°. The domains account for the different rankings: at 10 mm on axis the full-field fractions are 14.70/4.61%, but local fractions are 2.03/2.62%. Tangential widths are larger for CAMM at every sampled location.

### S2. Measured responses and region definitions

#### S2.1. Reflector measurements and eligibility

Five C5-2v cross-target records N1–N5 contain 20 buffer frames each, with 128 receive channels and 24 transmissions per frame. The sampling rate is 13.8889 MHz and reconstruction band 2.15–4.99 MHz. DAS and CAMM share RF, receive taper, band, and source clock; CAMM mode and phase-screen settings are fixed. TGC control points are $[0, 1023, 1023, 1023, 1023, 1023, 1023, 1023]$.

Scanner-reference images determine isolated-target coordinates independently of the paired reconstructions. The vertical column and rows near 19 and 39 mm give 18, 18, 20, 18, and 20 distinct windows in N1–N5; intersections count once and close-spaced clusters are excluded. Widths use independent RF reconstruction on $4 \times 6$-mm radial–tangential grids at 25-$\mu$m spacing. Half-amplitude crossings are linearly interpolated through each method's peak. A frame enters both methods' width statistics only when both peaks are interior and all four widths are bracketed. This excludes 18/1880 frame–window pairs, all in N5, leaving 16–20 paired frames per target. These widths characterize the reconstructed target response.

For reference target $(r_j, \theta_j)$, $s = r_j(\theta - \theta_j)$ defines inner $\mathcal{N}_j = \{|r - r_j| \leq 1 \text{ mm}, |s| \leq 3 \text{ mm}\}$ and outer $W_j = \{|r - r_j| \leq 4 \text{ mm}, |s| \leq 6 \text{ mm}\}$. Each polar cell contributes its intersection area with $W_j$ or $W_j \setminus \mathcal{N}_j$, holding its squared complex magnitude constant. The same masks and $512 \times 513$ full-field images are used for both methods, without background subtraction or interpolation; all 20 frames enter this integral. A complete $W_j$ must lie inside the field and contain no distinct scanner-reference peak within $W_j$ enlarged by 1 mm in both directions. Detections within 1 mm of the center are one candidate. This excludes N1/P13 and N3/P12 from energy alone.

**Table S2:** Measured local energy by record: medians of per-window framewise medians. Counts describe target windows within the five recorded acquisitions.

| Record | Width windows | Energy windows | CAMM lower | DAS (%) | CAMM (%) |
|---|---|---|---|---|---|
| N1 | 18 | 17 | 17 | 26.04 | 23.27 |
| N2 | 18 | 18 | 18 | 27.18 | 25.11 |
| N3 | 20 | 19 | 19 | 28.49 | 23.99 |
| N4 | 18 | 18 | 18 | 27.62 | 21.42 |
| N5 | 20 | 20 | 20 | 34.05 | 31.13 |
| All | 94 | 92 | 92 | 28.47 | 25.28 |

The median across windows of the framewise median CAMM-minus-DAS difference is $-2.85$ percentage points; it differs from subtracting the separately aggregated columns. Scaling both inner half-widths by 0.8 or 1.2 preserves the paired direction at all 92 windows. CAMM tangential width is larger at 91/94 targets. At peripheral N3/P29, asymmetric broadening accompanies median paired peak distance 0.665 mm and maximum 1.252 mm. These distances quantify the separation of the paired reconstructed peaks.

An acquisition-quality check flags 53 target echo windows with samples at $\pm 16384$ within $\pm 0.75$ $\mu$s of predicted transmit–receive arrivals, considering nonzero-weight receivers and all frames. Flagged targets remain in the eligible comparisons. The same directions persist in all 41 unflagged width windows and 39 unflagged energy windows; the latter median paired energy difference is $-2.59$ percentage points.

#### S2.2. Fixed cyst and carotid regions

Table S3 specifies the current masks in apex-based Cartesian coordinates. C1/C2 masks are fixed from the scanner-reference image. The carotid longitudinal and transverse views are from the same participant, the acquiring author. Within each record, masks are identical across methods and all 20 buffer frames. Repeated frames describe within-record variation.

**Table S3:** Fixed contrast-evaluation masks. All coordinates and radii are in mm.

| Region | Interior mask | Background masks |
|---|---|---|
| C1 | Disk: center $(-12.8, 28.7)$, radius 1.0 | Disks: $(-17.8, 28.7)$, $(-7.8, 28.7)$, radius 1.4 |
| C2 | Disk: center $(-23.0, 48.0)$, radius 1.7 | Disks: $(-30, 48)$, $(-16, 48)$, radius 1.8 |
| Longitudinal lumen | Polygon: $(13, 12.8)$, $(23, 10.8)$, $(23, 14.3)$, $(18, 15.8)$, $(13, 16.3)$; area 37.5 mm$^2$ | Disks: $(6, 22)$ and $(15, 22)$, radius 1.8 |
| Transverse lumen | Disk: center $(-2, 15.5)$, radius 1.8 | Disks: $(-11, 15.5)$ and $(7, 15.5)$, radius 1.8 |

Polar-cell areas weight uncompressed envelope means, variances, and histograms. Each method's envelope is divided by its own background mean. For each paired frame, 64 common bins span the pooled normalized interior and background samples from both methods. Measured CR is in dB; CNR is linear and gCNR dimensionless. C1/C2 radial depths are 29.7/50.7 mm and angles $-9.3°/-13.3°$, differing from Cartesian depths away from the axis.

The nominal longitudinal ROI gives more negative CR and higher CNR and gCNR for CAMM in all 20 frames. Shifting paired masks by $\pm 0.5$ mm in either coordinate or reducing the interior polygon to 90% of its linear dimensions preserves the signs of median paired differences. Nominal masks and main-text measurements are unchanged.

#### S2.3. Shallow candidate inclusion C0

The phantom layout locates C0 above the larger inclusions, although its boundary is indistinct in scanner images. A layout-predicted center is refined to minimum scanner intensity after 0.3-mm Gaussian smoothing in a fixed $\pm 1$-mm box; selection uses the scanner-reference image. The nominal interior disk has radius 0.4 mm; background disks of radius 0.65 mm lie 2.5 mm to either side at the same depth. Independent 25-$\mu$m RF reconstructions resolve these evaluation masks; their dimensions specify the sampled image regions.

**Table S4:** C0 nominal medians over 20 frames. Paired metric entries are DAS/CAMM; CNR is linear.

| Record | $x$ (mm) | $z$ (mm) | CR (dB) | CNR | gCNR |
|---|---|---|---|---|---|
| N1 | -12.59 | 9.49 | -4.38/-5.00 | 0.610/0.721 | 0.393/0.463 |
| N2 | -4.83 | 9.06 | 1.56/-2.39 | 0.311/0.328 | 0.255/0.250 |
| N3 | 12.43 | 6.69 | -2.77/-4.97 | 0.431/0.831 | 0.275/0.520 |
| N4 | 21.05 | 6.69 | -0.75/0.04 | 0.126/0.088 | 0.299/0.262 |
| N5 | -8.71 | 7.55 | 2.35/-4.78 | 0.413/0.653 | 0.274/0.386 |

Sensitivity checks use 16, 32, and 64 bins, interior radii 0.3, 0.4, and 0.5 mm, and separate $\pm 0.2$-mm center shifts in $x$ or $z$ with fixed backgrounds. Paired CNR and gCNR difference signs reverse in N1, N2, N4, and N5; N3 stays positive. All five echo windows overlap limit-valued RF samples under S2.1's rule. These measurements show record- and ROI-dependent small-region contrast.

### S3. C5-2v cyst studies

#### S3.1. Spatial configurations and size controls

One $20 \times 20$-mm tangent–radial template contains 24 000 uniformly positioned scatterers with independent Gaussian amplitudes, zeroed inside a radius-$a$ circle. Rotation and translation reuse the template without redrawing. Sixteen radius-4-mm cases cover $d = \{10, 15, 30, 50\}$ mm and $\theta = \{0, -5, -10, -20\}°$; two radius-2-mm controls at 30 mm complete the 18 configurations. Each row describes one paired realization.

The model uses 128 finite elements, 64 elevation subdivisions per element, the recorded 24-event sequence, and an 83.3333-MHz internal pulse clock followed by decimation to 13.8889 MHz. Both methods use the same RF and $512 \times 513$ grid. The interior ROI radius is $0.8a$ and background annulus $1.3a$–$1.8a$. Uncompressed envelopes use polar-cell area weights; 64 shared histogram bins span pooled normalized interior/background samples after division by each method's own background mean. Simulated CR and CNR are reported in dB.

**Table S5:** All 18 spatial configurations. More negative CR and higher CNR/gCNR indicate better target–background separation.

| Configuration | | | DAS | | | CAMM | | |
|---|---|---|---|---|---|---|---|---|
| $d$ (mm) | $\theta$ (°) | $a$ (mm) | CR (dB) | CNR (dB) | gCNR | CR (dB) | CNR (dB) | gCNR |
| 10 | 0 | 4 | -24.771 | 2.366 | 0.949 | -26.240 | 3.384 | 0.955 |
| 10 | -5 | 4 | -24.080 | 2.158 | 0.927 | -26.007 | 3.229 | 0.945 |
| 10 | -10 | 4 | -23.276 | 1.546 | 0.903 | -25.480 | 2.408 | 0.938 |
| 10 | -20 | 4 | -18.975 | -0.639 | 0.709 | -23.204 | -0.530 | 0.757 |
| 15 | 0 | 4 | -24.174 | 3.062 | 0.950 | -24.652 | 3.548 | 0.957 |
| 15 | -5 | 4 | -23.862 | 3.041 | 0.941 | -24.141 | 3.554 | 0.950 |
| 15 | -10 | 4 | -23.668 | 2.783 | 0.950 | -25.093 | 3.398 | 0.969 |
| 15 | -20 | 4 | -22.642 | 1.398 | 0.901 | -27.802 | 1.899 | 0.957 |
| 30 | 0 | 4 | -23.665 | 1.395 | 0.930 | -23.343 | 1.449 | 0.943 |
| 30 | -5 | 4 | -22.703 | 1.358 | 0.937 | -22.506 | 1.441 | 0.933 |
| 30 | -10 | 4 | -23.898 | 1.589 | 0.934 | -24.110 | 1.606 | 0.943 |
| 30 | -20 | 4 | -24.559 | 1.544 | 0.948 | -25.966 | 1.483 | 0.952 |
| 50 | 0 | 4 | -18.996 | 4.493 | 0.944 | -18.608 | 4.452 | 0.932 |
| 50 | -5 | 4 | -21.058 | 4.724 | 0.953 | -20.438 | 4.611 | 0.938 |
| 50 | -10 | 4 | -21.854 | 4.826 | 0.954 | -22.085 | 4.837 | 0.959 |
| 50 | -20 | 4 | -23.314 | 4.786 | 0.964 | -23.636 | 4.700 | 0.956 |
| 30 | 0 | 2 | -18.484 | 2.792 | 0.866 | -18.243 | 2.515 | 0.885 |
| 30 | -20 | 2 | -17.260 | 3.317 | 0.877 | -19.509 | 2.299 | 0.893 |

All eight 10–15-mm cases favor CAMM in all measures. Among the eight radius-4-mm cases at 30–50 mm, two favor CAMM, two favor DAS, and four have mixed rankings. At 30 mm/$-5°$, for example, CAMM-minus-DAS differences are $+0.197$ dB in CR, $+0.083$ dB in CNR, and $-0.004$ in gCNR. The radius-2-mm controls also have mixed rankings (Fig. S1).

The pulse construction repeats each excitation-state level for its recorded integer duration, reproducing the nonzero stored three-level waveform. Its sample count and duration imply a 250-MHz reference clock. Convolution with the recorded one-way impulse response agrees with the stored one-way waveform within $10^{-10}$ relative $L_2$ error. This reference underlies the lower-clock comparison in the main manuscript.

#### S3.2. On-axis cysts with varying realizations

Each realization contains three 20 × 20-mm planar patches at $(x, z) = (0, 25), (0, 60), (0, 100)$ mm, each with 24 000 scatterers and a radius-4-mm anechoic circle. Five independently seeded Gaussian-amplitude realizations use the same acquisition and ROI settings. At each depth, five distinct position seeds are used; some translated relative-position catalogues recur across depth–realization combinations. Each depth is summarized using its five realizations. Table S6 gives means and sample standard deviations; CR and CNR are averaged in dB.

**Table S6:** On-axis contrast across five realizations per depth: mean ± sample SD.

| $z$ (mm) | Method | CR (dB) | CNR (dB) | gCNR |
|---|---|---|---|---|
| 25 | DAS | $-22.05 \pm 0.44$ | $3.84 \pm 0.43$ | $0.955 \pm 0.008$ |
| | CAMM | $-21.17 \pm 0.43$ | $4.29 \pm 0.29$ | $0.949 \pm 0.006$ |
| 60 | DAS | $-18.90 \pm 0.35$ | $4.19 \pm 0.16$ | $0.928 \pm 0.005$ |
| | CAMM | $-18.31 \pm 0.30$ | $4.05 \pm 0.22$ | $0.922 \pm 0.016$ |
| 100 | DAS | $-20.64 \pm 0.68$ | $4.70 \pm 0.41$ | $0.944 \pm 0.012$ |
| | CAMM | $-20.12 \pm 0.74$ | $4.64 \pm 0.42$ | $0.940 \pm 0.008$ |

Mean CAMM gCNR is slightly lower at all three depths; mean CNR is higher at 25 mm and slightly lower at 60 and 100 mm. This comparison and the shared-template scan assess variability at fixed positions and spatial changes for one matched scattering realization, respectively.

#### Size-control images from the spatial study (S3.1)

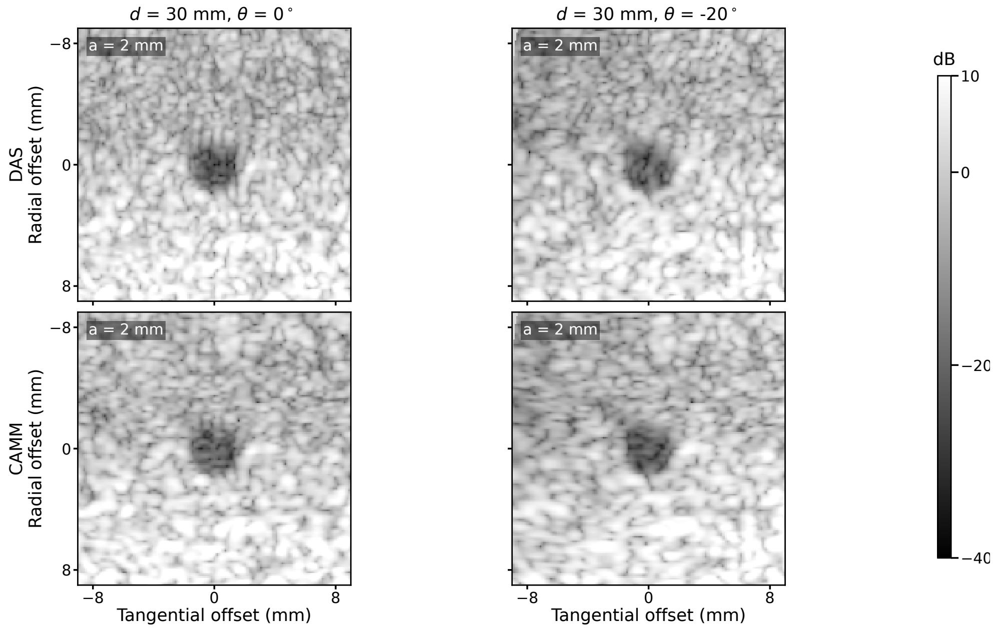


**Figure S1:** Radius-2-mm controls at 30 mm: on axis and at $-20°$, DAS above CAMM. Windows span 18 × 18 mm tangentially/radially; display levels are −40 to 10 dB relative to each method's area-weighted background mean. Display resampling is separate from the original polar-grid metrics in Table S5.

### S4. Computational measurements

An Intel Core i7-13700KF workstation (16 cores, 24 threads), 64 GiB RAM, and NVIDIA RTX 4090 (24 GiB) runs Windows 11 Pro, Python 3.11.4, and CuPy 13.6.0, separately from the acquisition console. Host latency starts with contiguous float32 RF in host memory and ends with the complex64 image returned to host memory, including transfers, RF transforms, reconstruction, and synchronization. File loading and B-mode display are excluded. Device latency uses CUDA events; preparation includes plan construction, static transfers, and warm-up. Two warm-ups precede eight alternating ABBA/BAAB blocks, giving 16 observations per method; host intervals are 10th–90th percentiles.

**Table S7:** One $3200 \times 128 \times 24$ recorded C5-2v frame on three grids. Host intervals are 10th–90th percentiles; other times are medians. Storage is persistent device arrays, including reusable buffers.

| Grid | Method | Host (ms) | Device (ms) | Prep. (s) | Storage (GiB) |
|---|---|---|---|---|---|
| $256 \times 257$ | DAS | 17.83 [17.00–19.54] | 12.50 | 2.47 | 0.15 |
| | CAMM dense | 48.10 [47.89–48.77] | 42.97 | 40.61 | 1.36 |
| | CAMM mixed | 37.32 [36.86–38.10] | 32.17 | 41.80 | 1.39 |
| $512 \times 513$ | DAS | 29.79 [29.02–30.60] | 23.72 | 7.40 | 0.26 |
| | CAMM dense | 85.30 [84.46–86.14] | 79.75 | 53.62 | 2.02 |
| | CAMM mixed | 51.87 [51.51–52.76] | 46.38 | 55.69 | 2.05 |
| $1024 \times 1025$ | DAS | 62.07 [60.96–62.81] | 54.48 | 24.95 | 0.71 |
| | CAMM dense | 183.10 [182.34–184.03] | 175.67 | 99.15 | 4.59 |
| | CAMM mixed | 120.71 [120.28–121.62] | 113.47 | 107.37 | 5.65 |

The physical field (radial depths 2.157–120.784 mm), RF, band, and prepared Stolt samples are shared across backend comparisons. The mixed policy selects low-rank radial products or NUFFT: all 23 intervals use low rank at $256 \times 257$ and $512 \times 513$; 14 use low rank and nine use NUFFT at $1024 \times 1025$. Relative spectral- and Frobenius-norm matrix tolerances are $10^{-5}$ for low rank; the NUFFT matrix check uses $2 \times 10^{-5}$. Maximum low-rank matrix errors at $512 \times 513$ are $9.94 \times 10^{-6}$ and $1.28 \times 10^{-6}$, respectively.

At that grid, low-rank evaluation reduces CAMM host latency from 85.30 to 51.87 ms. Its relative complex-image difference from dense phase-screen CAMM is $5.38 \times 10^{-7}$ at the original gain and phase. DAS remains faster and uses less persistent storage at all three grids. Reusable plans amortize preparation across frames with fixed acquisition and processing settings; the host times retain per-frame computation.

A separate comparison selects 4, 8, 12, and 24 approximately uniformly spaced transmissions, including both angular endpoints, at $512 \times 513$ with unchanged receivers, band, tolerances, and unit transmit weights. From 4 to 24 events, DAS/CAMM host times increase from 5.76/10.21 to 29.91/52.34 ms, and peak active CuPy-pool allocations from 184/762 to 298/2667 MiB. Table S7 reports a separate set of timing runs. Allocation peaks are measured after warm-up in separate method-only processes, including active plans and temporaries. They differ from persistent-array storage and exclude CUDA-context and vendor allocations outside the CuPy pool.
